# Pyridoxal 5’-phosphate supplementation modulates the heterologous expression and activity of a PLP dependent model protein in *E.coli*

**DOI:** 10.1101/2021.07.25.453669

**Authors:** Vijay Kumar Saxena, GV Vedamurthy, Raghvendar Singh

**Affiliations:** Molecular Physiology Laboratory, Division of Physiology and Biochemistry, Central Sheep and Wool Research Institute, ICAR-CSWRI, Avikanagar, Rajasthan- 304501, India; Livestock Research Centre, Southern Regional Station-, National Dairy Research Institute, ICAR-NDRI (SRS), Bengaluru, Karnataka-560030, India

**Keywords:** Pyridoxal 5’-phosphate, PLP, Protein expression, Cofactor, *Escherichia coli*, Fluorescence Quenching

## Abstract

PLP is a biologically active form of Vitamin B6 and is required for carbohydrates, amino acids and fatty acid metabolism. Many of the PLP dependent proteins are important drug targets and effector molecules, and thus, their heterologous overexpression is of industrial importance and has commercial value. We have predicted the docking site of PLP on O-acetyl serine sulfhydrylase protein (OASS) of *H*.*contortus* and determined that lysine-47 is very important for the binding of PLP in the enzyme pocket. We have used this protein as a model protein for testing the effect of PLP on the expression of PLP dependent proteins by *E*.*coli*. We have tested the effect of supplementation of PLP in the media on the expression of PLP dependent model protein in *E*.*coli*. Soluble recombinant protein could be purified from each of the culture vials grown with variable amount of PLP [0 mM (Group I), 0.01mM (Group II), 0.025mM (Group III), 0.05mM (Group IV) and 0.1mM (Group V)]. There was approximately 4.2%, 7.2%, 10.5% and 18% increase in purified protein yield in Group II, III, IV and V, respectively, in comparison to group I. We studied the relative incorporation of PLP into the purified protein by scanning the changes in internal fluorescence of purified proteins. There was a significant quenching of tryptophan fluorescence emission in groups II, III, IV and V compared to group I (Purified protein without PLP addition). There was a linear increase in the activity of protein purified from cultures of group I to group V. This was due to greater availability of PLP, thus, allowing higher incorporation of the cofactor in the apoenzyme to form holoenzyme complexes. PLP is not known to be directly imported into *E*.*coli*. We could find a PLP concentration-dependent increase in expression and catalytic activity of the enzyme signifying the probable transport of PLP across the membrane. The mechanism of transport of PLP in the light of the current experiment is still unknown and should be a subject of future studies.

## 1.0 Introduction

*Escherichia coli* has become the organism of choice to produce recombinant proteins and is used as a cell factory, being the most straightforward system for heterologous protein expression. In the last few decades, the development of recombinant DNA technology has considerably offset the need for the bulk of animal and plant tissues material to purify small amounts of a given protein. This unique ability to express and purify the desired recombinant proteins in large quantities has given us the leverage for the fast scaling up of proteins for their biochemical characterization, industrial and therapeutic utilization, and the development of commercial goods. The high yield of the protein and its solubility serve as the lynchpin for the commercial viability of the industrial scaling-up processes. In recent years, numerous new strains, vectors, and expression tags have been developed to overcome this system’s various limitations like codon biases, inclusion body formation, toxicity, mRNA instability, protein inactivity, and lack of post-translational modification **[1]**.

PLP is a biologically active form of Vitamin B6 and is essentially required for carbohydrates, amino acids, and fatty acid metabolism **[2]**. Enzymes utilizing PLP as cofactors are among the most abundant enzymes present in nearly all organisms. They catalyze many important reactions like racemization, decarboxylation, transamination, β, and γ- elimination reactions **[3,4]**. The unique ability of PLP to form covalent linkages (internal aldimine linkage with lysine residue of protein) allows them to function as electrophilic catalysts, stabilizing the different types of carbanionic reaction intermediates **[5,6,7]**.

The functional diversity of PLP-dependent enzymes can be readily appreciated from the fact that more than 184 different enzymatic activities **[8]** are PLP dependent, making up to ∼4% of all classified activities **[9]**. Since many of them are critical regulatory enzymes, they have been employed as potential drug targets **[10,11,12]**. Thus, any intervention which augments the heterologous protein expression of PLP-dependent proteins in *E*.*coli* would undoubtedly be a lynchpin in scaling up drug and target protein production.

Pyridoxal phosphate is one of the six interconvertible Vit B6 species, i.e., pyridoxal (PL), pyridoxine (PN), and pyridoxamine (PM) and their phosphate forms. They are also known as vitamers of pyridoxal phosphate. Only PLP and PMP (in few cases only) are the active catalytic forms of the vitamins. Vertebrates lack the enzymatic machinery for the synthesis of Vitamin B6. They are thus dependent upon the salvage pathway for recycling the alternate vitamer forms acquired from the diet **[13]**. Only microorganisms and plants could synthesize the vitamin *de novo* using either DXP dependent and DXP independent pathways **[14]**. PLP is known to be a membrane-impermeable molecule **[15,16]**, and excess of PLP was recently identified to be toxic to *E*.*coli* **[15]**.

There have been many studies on many PLP dependent proteins where PLP has assisted in the folding as well as structural stabilization, although the specific mechanism differs depending upon the particular protein and local microenvironment **[17,18]**. We hypothesized that if we can make available the PLP molecules during induction stage of protein expression, we can augment the soluble protein yields of PLP dependent proteins. To test this hypothesis, we took a PLP dependent O-acetylserine sulphadrylase protein of *Haemonchus contortus*, which was earlier characterized in our laboratory (GenBank MN733957) [**19**], as a model protein. Importantly, we also wished to reinvestigate whether the PLP, if added to media, could permeabilize the *E*.*coli* using our unique PLP dependent model protein. The protein model is unique as it has a single tryptophan residue (structurally significant) just next to a lysine, which could be used for probing comparative structural stabilization on the binding of PLP. This is done by monitoring the changes in the internal fluorescence of protein. We could ascertain by performing docking studies that lysine is very important for the binding of PLP as it contributes to the hydrogen bonding interaction with PLP. We expressed and induced OASS protein production at serially increasing concentrations of PLP supplemented in media. We have tried to probe the effect of the intervention on the expression, purification, protein folding, structural stabilization, and enzymatic activity of the model protein. The idea behind the study was that if the PLP is entirely impermeable, there should not be any significant difference in the above-studied parameters among different groups.

## 2.0 Material and Methods

### 2.1 3-D homology modelling of the OASS protein and prediction of the docking site of the cofactor PLP of the enzyme

Online version of the Phyre-2 (Protein Homology/analogy Recognition Engine V 2.0) web portal was used to assess the tentative secondary structure folding conformation of the protein. Homology modeling was done to determine the tentative structural folding taken by the receptor (entire CDS) in its native state. 3-D homology model of the CDS portion was prepared using (c3pc3A) as a template using the above software devised by Kelly et.al, 2015 [**20**] according to the standard methodology and validated as previously described [**21**]. Side chain modelling in Phyre-2 is based on fitting likely side chain rotamer conformations onto a fixed backbone. GalaxyDockWEB [**22**] was used to predict the 3D structures of protein-ligand complexes. With a given model structure of protein and a ligand structure, protein-ligand complex structures are generated by GalaxyDock protein-ligand docking program. The receptor is treated rigid by the program, and the ligand is treated flexible during docking simulation. The very purpose of modelling the docking site was to predict the critical amino acids and their conformational position with respect to bound pyridoxal L phosphate (PLP).

### 2.2 Expression of OASS at a variable concentration of PLP

An expression cassette for the production of recombinant OASS of *Haemonchus contortus* with a C-terminal hexa-his tag was cloned and produced in pET303 vector as part of another study (Saxena et al., 2021). The vector cassette was transformed in pLysS (BL21) strain of *E. coli* using a transformaid bacterial transformation kit (Thermofischer Scientific). Positive clones were verified by colony PCR and release of insert following double digestion with XbaI and XhoI. Five batches of cultures (50ml each) were inoculated in Luria-Bertani (LB) media containing ampicillin (100µg/ml), chloramphenicol (25µg/ml) at 37°C and variable concentrations of PLP (Sigma Aldrich) (0mM, 0.01mM, 0.025mM, 0.05mM and 0.1mM in batch cultures I,II,III,IV and V respectively) and induced by isopropy-ß D-thiogalactosidase (IPTG) at final concentration of 0.5mM for 4 hours.

### 2.3 Ni-NTA Affinity chromatography for the purification of proteins

The cultures were pelleted down at 4°C, and the pellets were dissolved in lysis buffer (200mM sodium phosphate and 300mM Sodium chloride, pH-8) with 1mg/mL Lysozyme and 1mM PMSF. The mixture was incubated on ice for 30 minutes, and the cells were lysed by cold shock 4-5 times with liq. N_2_ and 37 °C. Then, they were placed on ice for 5 minutes and centrifuged at 12,000 rpm for 30 minutes at 4°C. The supernatant was filtered by a 0.22µm filter, and the histidine-tagged fusion protein was purified from the filtered supernatant using the Ni-NTA agarose column (Thermo Fischer Scientific, USA). We took nearly 500μL of Ni-NTA beads for each culture batches, as we did not want the binding to be a limiting factor for any of our batch cultures, as we intended to investigate the relative protein yields. The bound proteins were eluted with 250mM imidazole concentration. The fractions of proteins from different culture batches were stored at -80°C for further use. The purity and relative concentration of purified recombinant OASS was analyzed by 12% sodium dodecyl sulphate polyacrylamide gel electrophoresis (SDS-PAGE) followed by Coomassie brilliant blue (R250) staining.

The concentration of protein purified out of different culture batches was purified by Pierce™ BCA Protein Assay Kit (Thermo Fischer Scientific). We also confirmed recombinant protein expression by immunoblotting using anti-His antibodies as standardized in earlier protocol from the laboratory **[23]**.

### 2.4 Comparative expression of protein at different induction conditions

A single colony was picked up from the transformed plate of bacteria with OASS expression cassette and was grown overnight in LB with 100μg ampicillin/mL at 37°C. The next day six cultures vials, each containing 10 mL of LB with 100μg ampicillin/ml, were inoculated with 50μL of overnight grown culture. The six culture vials were induced at variable PLP concentration as depicted in Table.1 when they reached an OD_600_ of 0.6 and were grown for 4h after induction.

**Table. 1.** Experimental group cultured at variable PLP concentrations.

| S.No | Culture groups | PLP concentrations used in culture (mM) | IPTG concentrations (mM) |
| --- | --- | --- | --- |
| 1. | Un-induced control | 0 | 0 |
| 2. | I | 0 | 1 |
| 3. | II | 0.01 | 1 |
| 4 | III | 0.025 | 1 |
| 5. | IV | 0.05 | 1 |
| 6. | V | 0.1 | 1 |

2mL of culture from each of the vials were precipitated at 12000rpm for 5min and re-suspended the cells in 60 μL of buffer (Tris buffer, pH-8.0), 20 μL of 10% SDS & 20 μL of 5X SDS loading dye and lysed by rapid vortexing and SDS PAGE was performed.

Another experiment was conducted as described in table 1, but cultures were grown at 16°C overnight after induction. The samples were prepared for SDS PAGE as already described above.

Densitometry was done by UVP image system software to analyze the relative changes in expression of the proteins among different groups.

### 2.5 Tryptophan quenching assay to assess the relative binding and structural stabilization of protein

Steady-state fluorescence measurements were performed with a Tecan Infinite M200 spectrophotometer (Tecan Life Sciences, Switzerland). Fluorescence emission of 5μg of the purified proteins dissolved in 50mM phosphate buffer (pH 7.4), was measured at 340 nm following excitation at 295 nm. Quenching of Trp fluorescence emission (due to variable association of PLP) was calculated as reduction of fluorescence intensity at λ_max_ 340nm, which suggest the formation of an association between the macromolecule and PLP due to changes in the Trp environment. We did not add PLP as an external quencher as we wanted to study the incorporation of PLP into protein while being expressed inside the *E*.*coli* cells.

### 2.6 Assessment of activity of purified OASS from different batch cultures (I-V)

20ug of each of the purified protein from each of the induced batch cultures was used to estimate O-acetyl serine sulphdrylase activity. The enzyme catalyzed the sulphdrylation of OAS using sodium sulphide to produce cysteine and acetate. The enzymatic reaction mixture of 100µL (200mM potassium phosphate buffer, 100mM DTT, 6.5mM O-acetyl serine) was added to the protein. Protein control was included, and the control for OAS was also included for each of the proteins. The mixture was incubated at 37°C for 5 minutes. Then, 5mM sodium sulfide was added, and reactions were incubated at 37°C for 30 minutes. After that, 15 µl of 20% TCA was added to stop the reaction, followed by its centrifugation at 12000*g for 5 minutes. Then, 100 µL of supernatant was added in 100µl of Acid Ninhydrin Reagent-2, freshly prepared. The reaction mixtures were boiled at 100°C for 10 minutes and cooled for 5 minutes on ice. Then, the absorbance was recorded at 560nm and analyzed with respect to control. L-cysteine (0–50 μmol) dissolved in acid-ninhydrin reagent was used as standard and treated similarly as described earlier to quantify the amount of cysteine formed.

### 2.7 Statistical Analysis

Results are presented as mean and standard deviation of the mean. One way ANOVA was used to assess the significance of differences in various parameters studied for all the *E*.*coli* groups. A probability level of < 0·05 was taken to indicate a statistically significant difference in the means between data pairs. Normalized Data were analyzed and graphs were plotted using the Graphpad Prism 8.

## 3.0 Results

### 3.1 Homology modelling and docking sites of PLP on OASS

Secondary structure analysis could demonstrate the predominance of alpha-helical secondary conformation, which was 45%, followed by β-sheet structure (15%) and random coil structure (9%) (**Fig. A**_**1**_ **Supplementary Figure**). 3-D homology model of the protein was prepared using c3pc3A (Uniprot sequence id) as a template using an online version of the Phyre-2 protein modelling software (**Fig A**_**2**_ **Supplementary Figure**). c3pc3A used as template was the full-length structure of cystathionine beta-synthase from drosophila2 in complex with aminoacrylate. The prepared model is quite good as 318 residues (95% of sequence) have been modelled with 100.0% confidence by the single highest scoring template. The template shows 35% identification (minimum requirement for homology modelling is 30%) with the target query sequence. Docking sites of PLP was predicted using Galaxyweb dock. **Fig 1a** suggests the location of PLP binding on the modelled structure of OASS. There are important interactions of amino-acids (Lys 47, Gly 182, Gly 184, Thr186, Cys187, Gly 225, Asn 297) on enzyme that help bind enzyme with the PLP and thus should be essentially crucial for activity of the enzyme **Fig1b**.

**Fig 1a.**
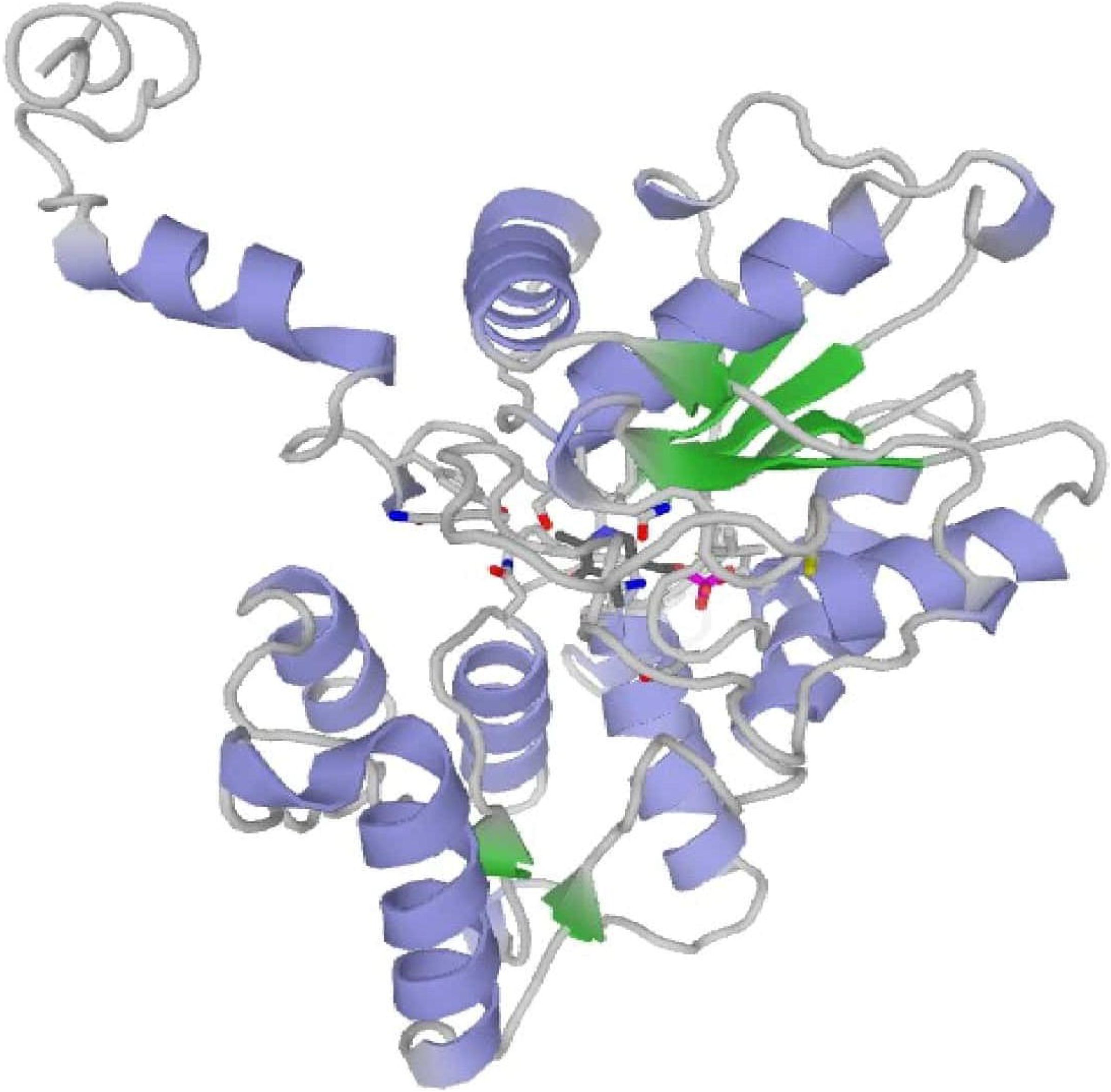
Docking sites prediction of PLP: Docking sites for PLP was predicted using Galaxyweb, dock location of PLP binding on the modeled structure of OASS.

**Fig 1b.**
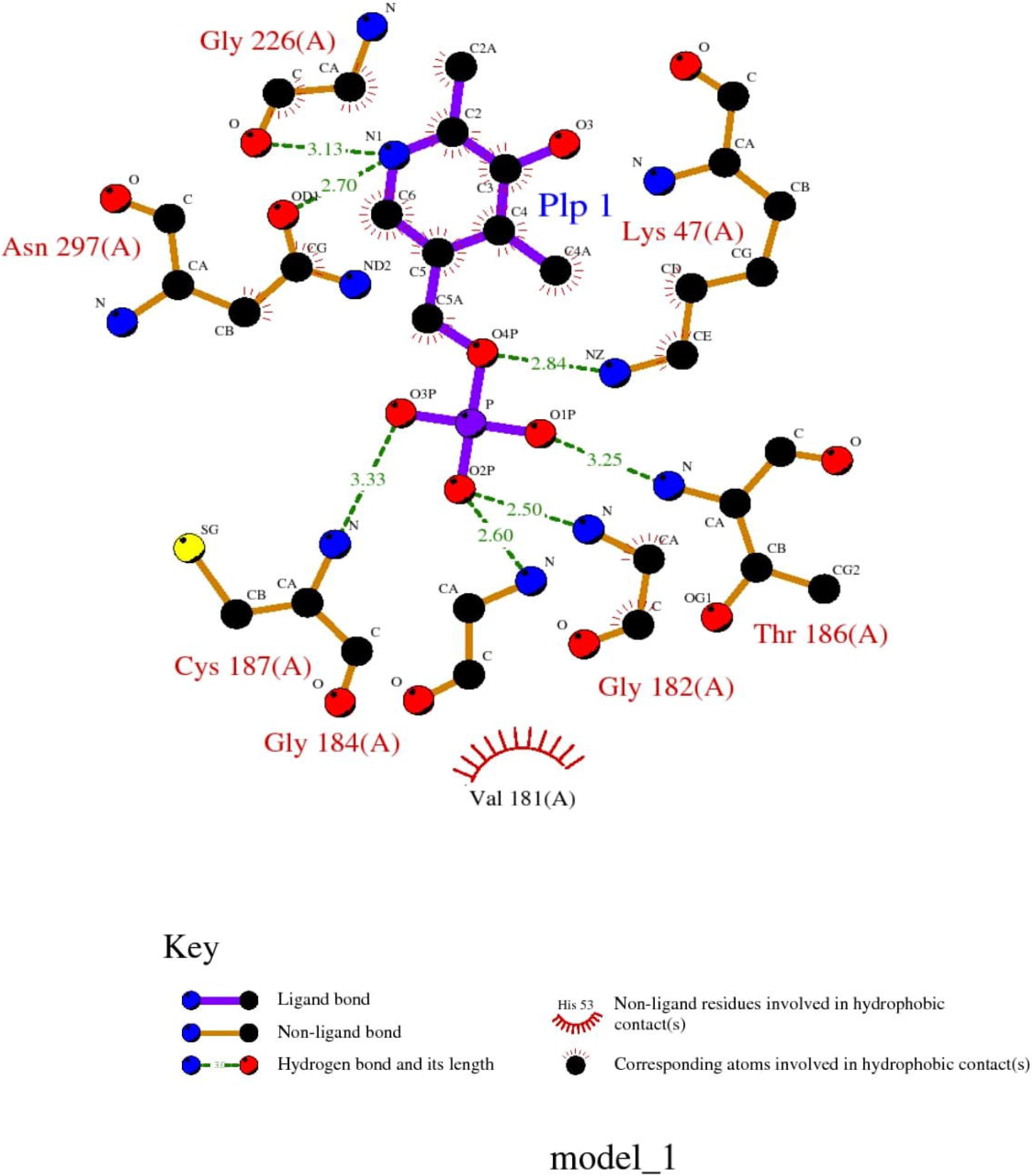
Important interactions of amino-acids (Lys 47, Gly 182, Gly 184, Thr186, Cys187, Gly 225, Asn 297: All marked in Red) on enzyme which help in the binding of enzyme with the PLP for activity of the enzyme was depicted in ball and stick model.

### 3.2 Targeted expression of model protein OASS

The expression cassette of the gene coding for the O-acetyl serine sulphdrylase of *H*.*contortus* was reverified by PCR and was transformed successfully into pLysS (BL21) strain of *E. coli*. The clones were found to be positive by colony PCR and further confirmed by restriction enzymes mediated release of the insert of 1003bp in size. (**Fig.A3 Supplementary Figure**). Induction with different concentrations of PLP did not negatively affect the growth profile of the pLysS (BL21) strain of *E. coli*. We did not find a significant difference among various groups in weight of pellet measured both at the time of induction by IPTG and 4h post-induction. We observed that even a concentration of 0.5mM PLP in media did not negatively affect the growth profile of pLysS (BL21) strain of *E. coli* with an expression cassette. Soluble recombinant protein of 38kDa in size could be purified from each of the culture vials grown with a variable amount of PLP [0 mM (Group I), 0.01mM (Group II), 0.025mM (Group III), 0.05mM (Group IV) and 0.1mM (Group V)]. There was approximately 4.2%, 7.2%, 10.5% and 18% increase in protein yield in Group II, III, IV and V, respectively, compared to group I. The respective protein yield from 50ml culture in each group is presented in **Fig.2a**. The purity of the protein was excellent, as it was purified as a single band in all the culture groups, and concentration was increasing from group II-V. As an equal volume (rather than concentration) of purified proteins was used for SDS PAGE, a very light band was obtained for protein from group I. It was found to increase in intensity from group II TO V (**Fig2b)**. The western blot (WB) using anti-His antibody could confirm recombinant OASS protein for all the culture groups (**Fig2c)**. The presence of protein at low concentration in group I was very well evident.

**Fig.2.**
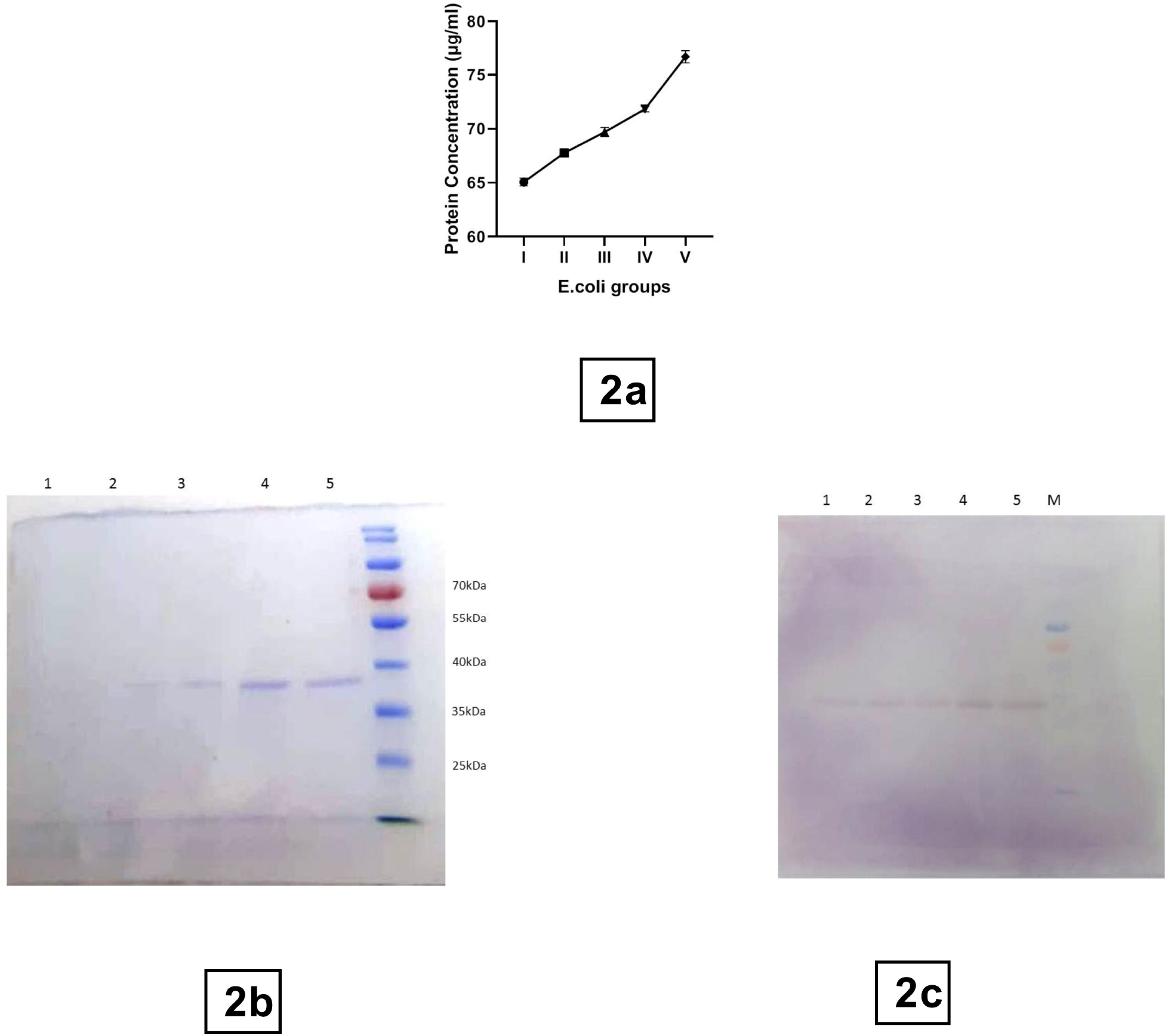
Expression of purified OASS out of different *E*.*coli* groups supplemented with variable PLP concentration. a) Concentration of purified OASS purified from 50ml culture volume of each of different *E*.*coli* groups (I to V), b) SDS PAGE analysis of equal volume of purified proteins clearly demonstrating the increase in expression of OASS from group I to group V, Lane 1-Lane 5 contains proteins purified from group I to V respectively, Lane M is molecular weight marker Page Ruler prestained marker (Thermofisher Scientific) c) Western blot confirmation of the protein and verification of increase in expression of OASS from group I to V. The lane depiction, markers and protein loading remains the same as Fig. 2b

### 3.3 Validation of enhanced expression using different induction conditions

The protein expression was increasing serially from Group I –V when we induced protein expression using 1mM IPTG at 37° C **(Fig 3a)**. We observed similar results when we induced the protein at low-temperature conditions **(Fig 3b)**. Best expression was found in the induced fraction treated with 0.1mM PLP (Group V) in the culture media for both the induction conditions. There was a significant impact of the addition of PLP on protein yields at all studied concentrations, for both induction at 37°C and low-temperature condition protocol. There was a non-significant difference in protein yields between different induction conditions (37°C and low-temperature conditions) at all studied concentrations of PLP (**Fig.4**).

**Fig. 3.**
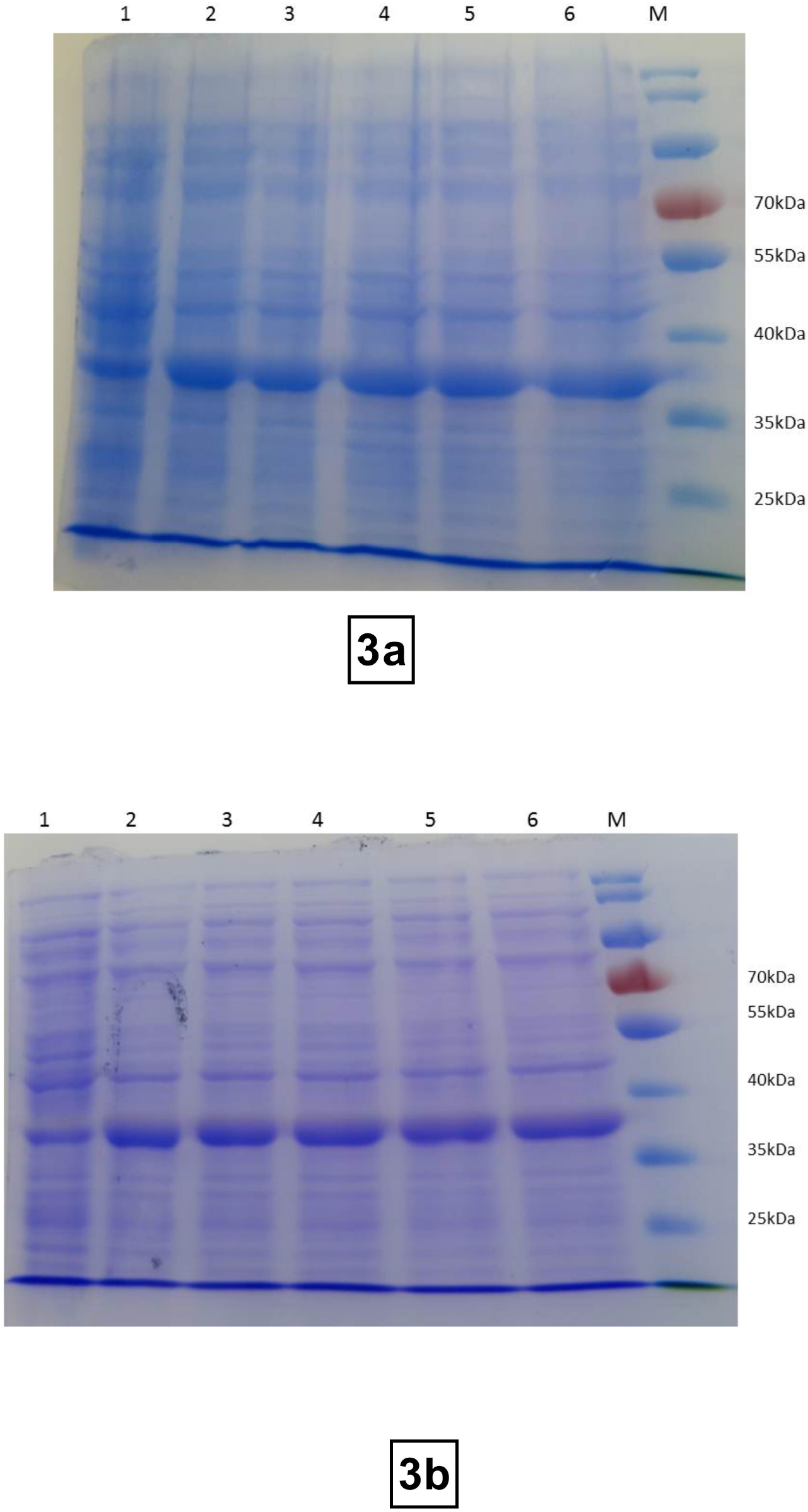
Comparison between normal induced proteins and proteins induced in presence of different concentrations of PLP a) OASS Protein induction at 37°C for 4h, Lane 1 is lysate of uninduced control, Lane 2-6 are induced protein from group I to V, b) OASS Protein induction at 16°C for overnight, Lane 1 is lysate of uninduced control, Lane 2-6 are induced protein from group I to V,

**Fig.4.**
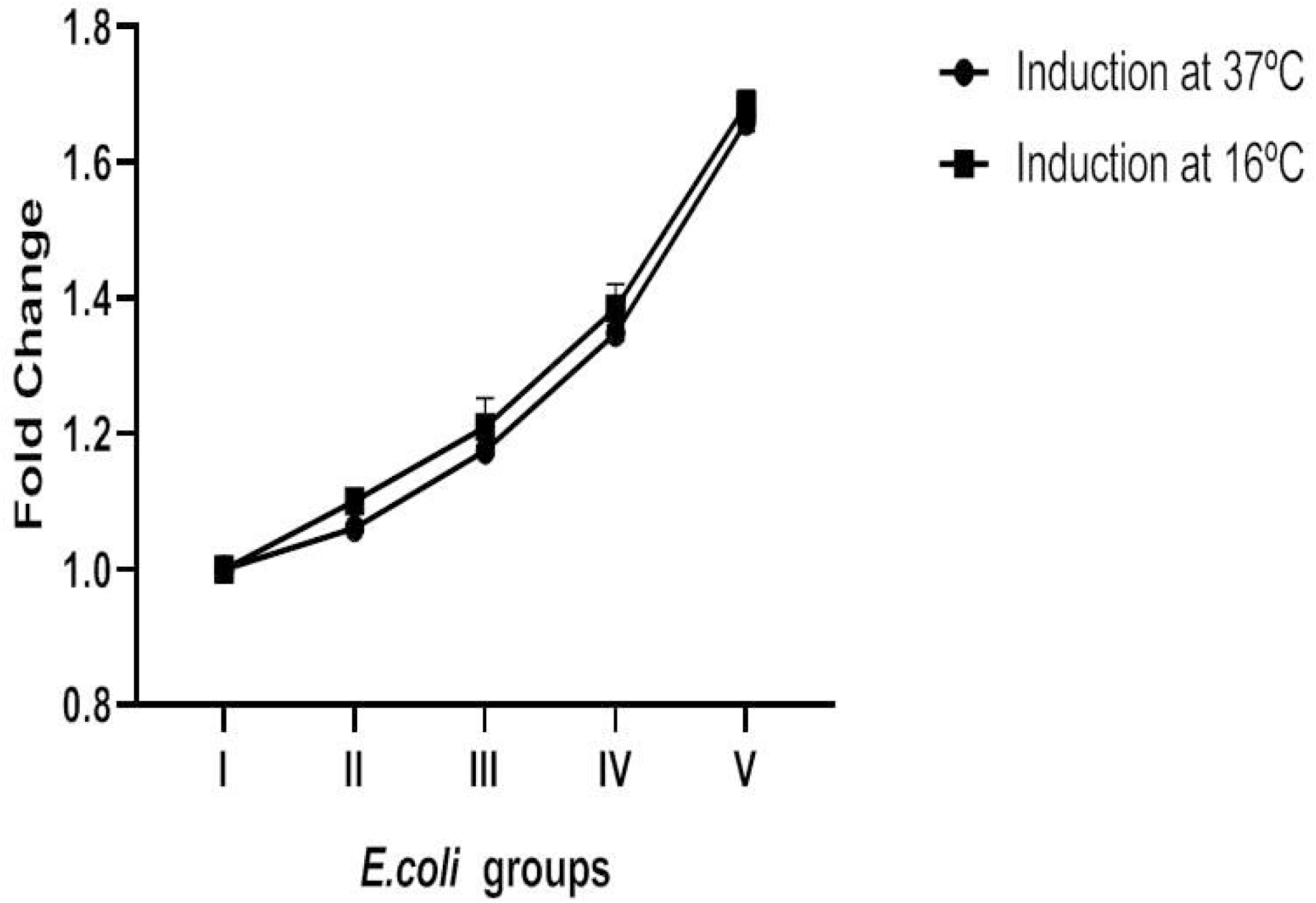
Analysis of relative expression of OASS protein in group II to V, in comparison to group I, by densitometric analysis under two different induction conditions.

### 3.4 Tryptophan quenching studies

There was a significant quenching of tryptophan fluorescence emission in groups II, III, IV and V compared to group I (Purified protein without PLP addition) (**Fig.5**). There was a significant difference in fluorescence emission among all groups.

**Fig.5.**
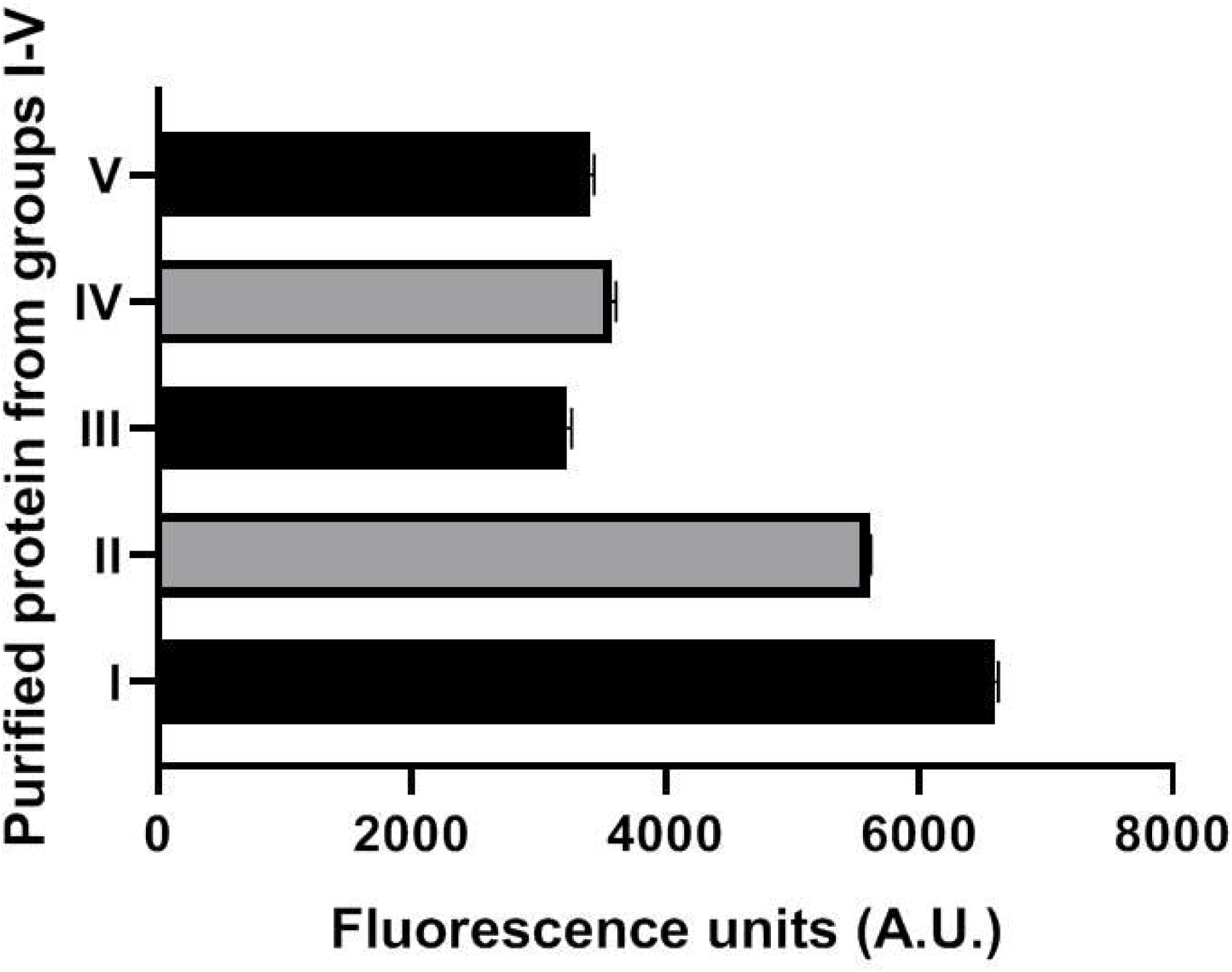
Quenching of internal fluorescence as observed by measuring Trp fluorescence emission at 340nm by equal concentration of OASS protein purified from different *E*.*coli* groups

### 3.5 Enzymatic activity of proteins isolated from different groups

The enzymatic activity was measured as the amount of cysteine produced for the reaction time of 30 minutes for all the protein groups taking an equal amount of protein for analysis. The enzymatic activity was serially increasing from group I to group V (**Fig.6**). There was a significant increase in enzyme activity in group IV and V compared to group I, signifying the best incorporation of the cofactor to the apoenzyme.

**Fig.6.**
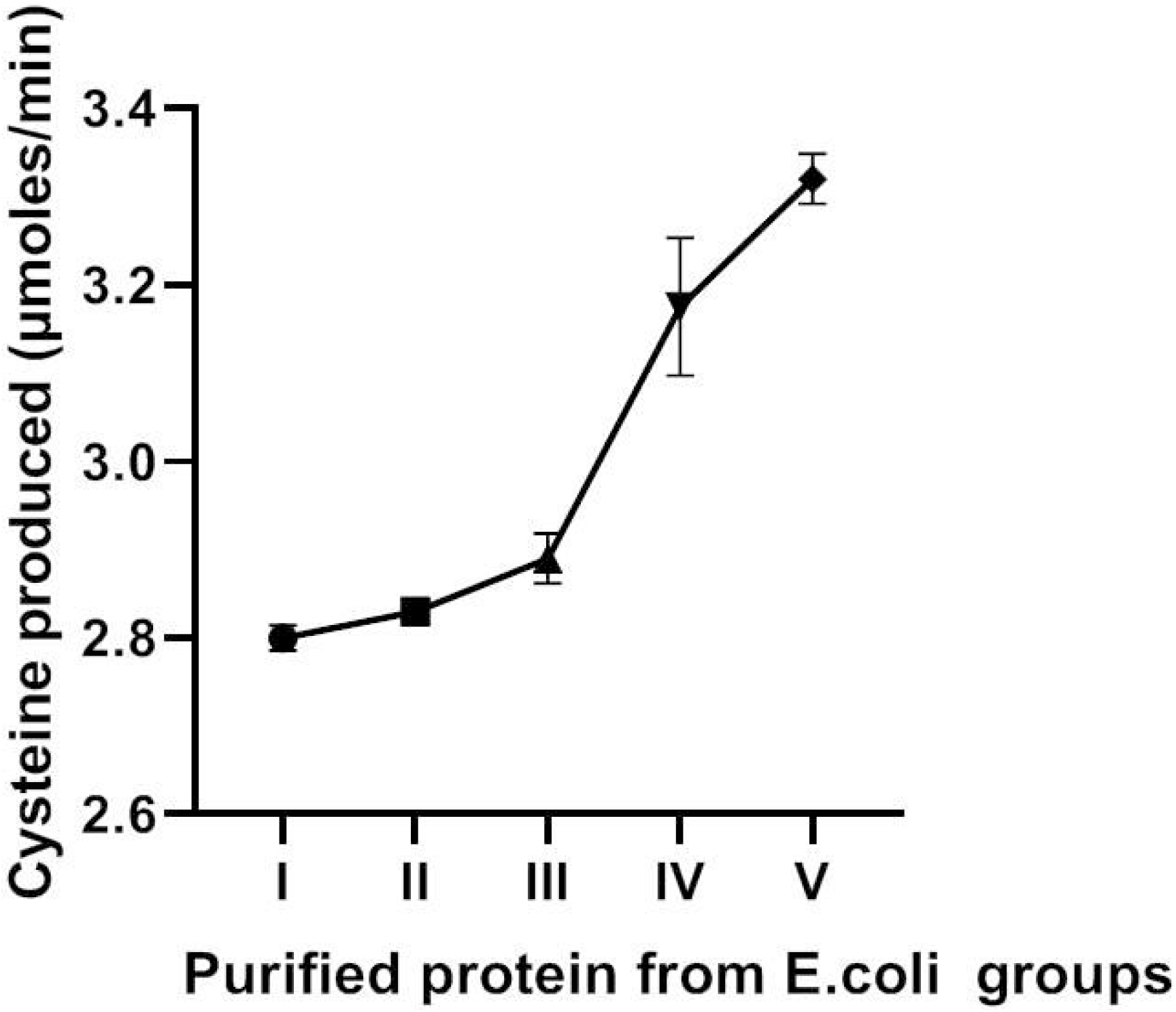
Enzyme activity of purified proteins from different *E*.*coli* groups measured as rate of cysteine produced per minute.

## 4.0 Discussion

### 4.1 Structural Characterization of OASS of *H*.*contortus*

The structural conformation of the OASS protein consisted primarily of alpha-helical (45%) and β-sheet conformation. Galaxy dock analysis for ligand mapping could identify PLP as its most important cofactor, which is in line with the basic properties of the cysteine synthases enzyme performing alanyl transfer using PLP as a co-factor. Lysine residue at 47^th^ place was found to be a much-conserved amino acid in Cysthathionine β synthase (CBS), ranging from bacteria to humans, which could be a crucial interactive amino acid PLP in the enzyme. This lysine could be involved with forming the internal aldimine complex, which is an essential mechanism of most of the PLP cofactored transamination enzymes. Kobylarz et al, 2016 [**24**] co-purified PLP with the enzyme SbnA, a pyridoxal 5-phosphate (PLP) dependent enzyme with homology to O-acetyl-L-serine sulfhydrylases. They observed that PLP was covalently bound to Lys47 at the base of the cleft by an internal Schiff base originating from the PLP psi-face. The enzyme used O-phospho serine (OPS) as the substrate. The PLP formed hydrogen bonds to Ser185, Thr186, Thr187, and Ser189 and three water molecules. Similarly, we observed that the conserved stretch of amino-acids VGSGGTC (181-187) in OASS protein required for the binding of PLP chiefly as a source of stabilizing the hydrogen bond formation with the PLP. Thus, the characterized OASS demonstrated the structural uniformity with PLP bound serine sulphdrylases and nearly utilized the same chemical bonding features and conformational arrangement of amino acids to bind PLP.

### 4.2 Effect of PLP on yield of model OASS protein

PLP dependent enzymes are important drug targets and effector molecules. Many of them are having diverse catalytic activities. PLP is a critical molecule for these proteins as it remains tightly bound to an internal lysine through the ε-amino group forming an internal aldimine complex and acts as an electron sink withdrawing electrons from the substrate **[25]**. Therefore, we hypothesized that if we supplement PLP in the media, it may positively affect the soluble expression and yield of the protein. In our experiment, we cultured pLysS (BL21) strain of *E. coli (*with a transformed expression cassette of OASS gene of *H*.*contortus)*, with serially increasing concentrations of PLP supplemented in media. We did not notice any toxic effect of PLP on the growth of the culture as there was no significant difference in mean OD_600_ at the stage of induction and the 4h post-induction stage. Sugimoto et al., 2017 **[15]** observed that excess PLP has a negative effect on the growth profile of the *E*.*coli*, which is due to the imbalance created in the amino-acid pool. His study concluded that excess PLP could hamper amino-acid acid metabolism, creating a shortage of some particular amino acids creating a growth defect. We may not have observed the toxicity at the studied concentrations of PLP, as the used concentrations may be optimal and non-toxic. The soluble protein expression increased with the increasing availability of PLP in the media. Since all the culture groups had been induced at a uniform stage and were given similar treatments, it suggests a positive effect of PLP on the soluble expression of the protein. The quantitative analysis of protein, western blot confirmation, corroborated that protein expression was increasing substantially from group I (where no PLP was added) to Group V (0.1mM PLP).

### 4.2 PLP, Protein folding and native state stabilization

More than 30% of the proteins require their cofactors for their biological activities. In case of heterologous expression in *E*.*coli*, where the microenvironment is considerably different, the availability and concentration of cofactor may have a very important role, especially if it’s helping in attaining the native stage of the protein. PLP has been studied to help attain the native stage for some of the PLP dependent proteins **[26,27]**. In addition to it, various earlier studies have demonstrated the role of PLP in the folding mechanism of some of the PLP dependent proteins **[26, 28,29,30]**. They have been studied to bind the unfolded state of the protein and serve as nucleation sites for faster folding of the protein, which could be a boon when looking at the scenario of heterologous protein expression. We induced protein expression at 37 °C for 4 h using a higher concentration of IPTG to increase the positive production pressure. We found that protein yield increased from group I to V, with an increase in PLP concentration in media.

We also studied low-temperature induction as it allows newly translated recombinant proteins more time for folding. We observed that supplementation of PLP in media increased the protein yields at different induction conditions (37 °C for 4 h and 16 °C for overnight) at all studied concentrations of PLP. There was a non-significant difference in protein yields when we studied the relative changes between different induction conditions (37 °C for 4 h and 16 °C for overnight), indicating that PLP, especially in the case of OASS, has more to do with native state stabilization rather than an active help in protein folding process. We tested this further with tryptophan quenching studies on the purified proteins from cultures induced at different concentrations of PLP. Proteins have an internal fluorescence due to the presence of aromatic amino acids, as they absorb light, promoting the transition of electrons to an excited state, resulting in the emission of light of a longer wavelength when electrons return to the ground state. The fluorescence of proteins (mainly due to tryptophan) can be selectively measured at an excitation wavelength of 295 nm. There is no absorption by tyrosine at this wavelength, and phenylalanine also does not contribute to it, owing to low quantum yield. Tryptophan fluorescence is highly sensitive to the polarity of its environment, and it changes its emission spectra in response to protein conformational transitions, subunit association, ligand binding, or denaturation, which affect the environment surrounding the indole ring **[31]**. Molecules like PLP have their fluorescence and can interact with the excited state of the tryptophan to yield an excited product that decays by a non-radiative pathway producing the dynamic quenching of tryptophan fluorescence. This quenching effect can be measured at the emission maxima of tryptophan, which is 340nm but may vary slightly (8-10nm) depending upon the protein. The protein OASS could be a perfect model protein for quenching studies with PLP, as it has a tryptophan residue at the very next to lysine in its primary structure. The suggested mechanism is that an increase in binding of PLP to the lysine-47 quenches the internal fluorescence of the protein due to excitation of tryptophan as it is in close proximity. We could obtain a significant quenching of tryptophan fluorescence in group II–V protein compared to control (Group I), which suggest an increase in conformational stabilization and greater incorporation of PLP in the protein, as PLP is an important quenching molecule. This also could justify the greater yield of protein expression and increase in protein activity with an increase in PLP supplementation in media.

### 4.3 Effect of PLP on functional enzyme activity

OASS catalyzes the sulphdrylation of O-acetyl serine to form cysteine as well as acetate. It’s an enzyme of de-novo pathway of cysteine synthesis recently characterized by our group in *H*.*contortus* [**19**]. The cysteine synthase activity of the enzyme could be accurately quantified by measuring the amount of produced cysteine, using a modified protocol of Gaitonde et.al, 1967 **[32]**. This method allows us to precisely measure amino-acid cysteine even in the presence of other alpha-amino acids. There was a linear increase in the activity of protein purified from cultures of group I to group V. This may be due to the greater availability of PLP, thereby leading to the higher incorporation of the cofactor in the apo-enzyme to form holoenzyme complex. Higher cofactor incorporation has been studied to increase enzyme activity, like carboxylic acid reductase (CAR), which has phosphopantetheine as a cofactor **[33]**. Sole expression of the enzyme carboxylic acid reductase (CAR) leads to poor activity, but it could be augmented by co-expression of phosphopantetheinyl transferase **[34]**. Phosphopantetheinyl transferase catalyzes the incorporation of phosphopantetheine into the CAR. Pyridoxine supplementation has been found to improve the activity of recombinant Glutamate Decarboxylase and the enzymatic production of gamma-aminobutyric acid (GABA) **[35]**. Pyridoxine is one of the vitameric forms of VitB_6_ and has been studied to be membrane permeable, while PLP is known to be membrane impermeable. Pyridoxine has to be salvaged to PLP to act as a cofactor for glutamate decarboxylase.

### 4.4 Does PLP internalization occurs in *E*.*coli*?

PLP is known to be membrane-impermeable **[16]**and thus is considered a form of the vitamin that can be very well entrapped inside cytoplasmic compartments in eukaryotes, where most PLP dependent enzymes are localized. Human cells have two active transport components for the transport of B6 vitamins; one is specific for unphosphorylated forms while the other is specific for phosphorylated forms **[36,37]**. Yamada et. al,1977 **[38]** performed the membrane uptake studies using radioactive vitamin derivatives in *E*.*coli. They* summarized that *Escherichia coli* takes up concentratively all three forms of nonphosphorylated vitamin B6 but does not take up the phosphorylated forms. Dempsey and Pachler, 1966 **[39]** also suggested that although PL, PM, and PN were imported, PLP is not imported into *E*.*coli* cells. Our innovative approach to PLP membrane transport measurements by catalytic amplification of the permeation signal through activation of a PLP-dependent enzyme indicates the probable transport of PLP across the membrane, but the exact transportation mechanism is to be studied in detail. The recombinant protein (OASS) was sensitive to the permeation signal (PLP) as it demonstrates the concentration increase in protein yield and enzyme activity (catalytic amplification). The significant quenching of internal fluorescence of purified OASS from PLP supplemented groups compared to control does signify a difference in the binding of PLP to the lysine. But there is a need to further confirm this by PLP internalization studies after labelling PLP with suitable fluorescent probes without compromising the native properties of the molecule.

As a result, we propose OASS of *H*.*contortus* as an excellent model system for linked permeation studies. PLP being non–toxic at studied concentrations, could be used for amplifying the expression of PLP linked proteins. They can be individually titrated for the increase in expression with different PLP concentrations. The approach developed by us gives a unique method for amplifying the expression and activity of PLP dependent proteins and enzymes by directly supplying the catalytically active cofactor molecules, i.e., PLP in media. PL, PM and PN (alternative forms of VitB_6,_ known to be a permeable form of vitamers), when supplied in media, may not be quickly incorporated, as they need to be converted to PLP by enzymes of salvage pathways. It may add to the metabolic burden as most metabolic machinery is diverted towards the heterologous protein expression in induced *E*.*coli* culture.

## Conclusion

The inclusion of PLP in growth media does bring about a dramatic increase in protein yield and enzymatic activity, which is probably by greater incorporation of PLP into the model PLP dependent protein. Based on our research findings, PLP appears to be getting imported into the *E*.*coli*, which is demonstrated by linked enzymatic machinery used in the study. Still, the actual confirmation of the process and mechanism of transport is to be studied in detail by intensive internalization studies. We report a new method for direct overexpression of PLP dependent proteins using PLP, rather than using other vitamers needed to be salvaged to PLP by *E*.*coli*.

## Abbreviations

PLP: Pyridoxal 5’-phosphate
PL: Pyridoxal
PM: Pyridoxamine
PN: Pyridoxine
OASS: O-acetylserine sulphdrylase
DXP: Deoxyxylulose 5-phosphate
CDS: Coding sequence

## Acknowledgement

The authors are thankful to Department of Science and Technology, Government of India, for providing the funding via ECR/2016/000135. Special thanks to the Indian Council of Agricultural Research (ICAR) and the Director, CSWRI, for providing the necessary support and infrastructure. We are also thankful to support provided by Shri Suryaprakash Sharma and Vishnu Sharma in the conduction of the studies.

## Author Credit Roles

**Vijay Kumar Saxena: Investigation**, Conceptualization; Methodology, Visualization, Validation, Data Curation, Writing Original Draft, Project Administration, Review & Editing: **Vedamurthy GV**–Methodology, Validation, Funding Acquisition, Review & Editing; **Raghvendar Singh:** Validation, Data curation, resources

## Figure Legends

**<u>Supplementary Fig.A</u>**_**<u>1</u>**_

Secondary structure conformation prediction using Phyre 2

**<u>Supplementary Fig.A</u>**<u>_**<u>2</u>**_</u>

3-D homology model: CS protein 3-D structure was prepared using c3pc3A_ (Uniprot sequence id) as a template using online version of the Phyre-2 protein modeling software.

**<u>Supplementary Fig.A</u>**_**<u>3</u>**_ a) Colony PCR of the obtained colonies confirming the presence of insert (1003bp) (L_1_-L_5_), L6 is 1kb MW marker generuler™ (Thermofisher Scientific) b) Restriction digestion confirmation of the cloning by the release of insert (10003bp), L1 is 1kb MW marker generuler™ Thermofisher Scientific, L2 is undigested control, L3 is digested plasmid with XbaI and XhoI.

## Notes

### Competing Interest Statement

The authors have declared no competing interest.

## References

1. Khow, O. and Suntrarachun, S., 2012. Strategies for production of active eukaryotic proteins in bacterial expression system. Asian Pacific journal of tropical biomedicine, 2(2), pp.159–162.

2. Mooney, S., Leuendorf, J.E., Hendrickson, C. and Hellmann, H., 2009. Vitamin B6: a long known compound of surprising complexity. Molecules, 14(1), pp.329–351.

3. Percudani, R. and Peracchi, A., 2009. The B6 database: a tool for the description and classification of vitamin B6-dependent enzymatic activities and of the corresponding protein families. BMC bioinformatics, 10(1), p.273.

4. Richts, B., Rosenberg, J. and Commichau, F.M., 2019. A Survey of Pyridoxal 5′-Phosphate-Dependent Proteins in the Gram-Positive Model Bacterium Bacillus subtilis. Frontiers in molecular biosciences, 6, pp.32.

5. Bettati, S., Benci, S., Campanini, B., Raboni, S., Chirico, G., Beretta, S., Schnackerz, K.D., Hazlett, T.L., Gratton, E. and Mozzarelli, A., 2000. Role of pyridoxal 5′-phosphate in the structural stabilization of O-acetylserine sulfhydrylase. Journal of Biological Chemistry, 275(51), pp.40244–40251.

6. John, R.A., 1995. Pyridoxal phosphate-dependent enzymes. Biochimica Et Biophysica Acta (BBA)-Protein Structure and Molecular Enzymology, 1248(2), pp.81–96.

7. Schneider G., Kack H. & Lindqvist Y. (2000) The manifold of vitamin B6 dependent enzymes. Structure Fold. Des., 8, R1–R6

8. Prunetti, L., El Yacoubi, B., Schiavon, C.R., Kirkpatrick, E., Huang, L., Bailly, M., El Badawi-Sidhu, M., Harrison, K., Gregory 3rd, J.F., Fiehn, O. and Hanson, A.D., 2016. Evidence that COG0325 proteins are involved in PLP homeostasis. Microbiology, 162(4), pp.694–706.

9. Percudani, R. and Peracchi, A., 2003. A genomic overview of pyridoxal-phosphate-dependent enzymes. EMBO reports, 4(9), pp.850–854.

10. Kleppner, S.R. and Tobin, A.J., 2001. GABA signalling: therapeutic targets for epilepsy, Parkinson’s disease and Huntington’s disease. Emerging Therapeutic Targets, 5(2), pp.219–239.

11. Snell, K. and Riches, D., 1989. Effects of a triazine antifolate (NSC 127755) on serine hydroxymethyltransferase in myeloma cells in culture. Cancer letters, 44(3), pp.217–220.

12. Wang, C.C., 1995. Molecular mechanisms and therapeutic approaches to the treatment of African trypanosomiasis. Annual Review of Pharmacology and Toxicology, 35(1), pp.93–127.

13. di Salvo, M.L., Contestabile, R. and Safo, M.K., 2011. Vitamin B6 salvage enzymes: mechanism, structure and regulation. Biochimica et Biophysica Acta (BBA)-Proteins and Proteomics, 1814(11), pp.1597–1608.

14. Fitzpatrick, T.B., Amrhein, N., Kappes, B., Macheroux, P., Tews, I. and Raschle, T., 2007. Two independent routes of de novo vitamin B6 biosynthesis: not that different after all. Biochemical Journal, 407(1), pp.1–13.

15. Sugimoto, R., Saito, N., Shimada, T., and Tanaka, K. (2017). Identification of YbhA as the pyridoxal 5′-phosphate (PLP) phosphatase in Escherichia coli: importance of PLP homeostasis on the bacterial growth. J. Gen. Appl. Microbiol. 63, 362–368.

16. Whittaker, J.W., 2016. Intracellular trafficking of the pyridoxal cofactor. Implications for health and metabolic disease. Archives of biochemistry and biophysics, 592, pp.20–26

17. Herold, M. and Leistler, B., 1992. Coenzyme binding of a folding intermediate of aspartate aminotransferase detected by HPLC fluorescence measurements. FEBS letters, 308(1), pp.26–29.

18. Bertoldi, M., Cellini, B., Laurents, D.V. and Borri Voltattorni, C., 2005. Folding pathway of the pyridoxal 5′-phosphate CS lyase MalY from Escherichia coli. Biochemical Journal, 389(3), pp.885–898.

19. Saxena V.K., Vedamurthy G.V., Swarnkar C.P., Kadam V., Onteru, S.K. Ahmad H., Singh R., De novo pathway is an active metabolic pathway of cysteine synthesis in Haemonchus contortus, Biochimie (2021), doi: https://doi.org/10.1016/j.biochi.2021.05.014

20. Kelley, L.A., Mezulis, S., Yates, C.M., Wass, M.N. and Sternberg, M.J., 2015. The Phyre2 web portal for protein modeling, prediction and analysis. Nature protocols, 10(6), p.845.

21. Shin, W.H. and Seok, C., 2012. GalaxyDock: protein–ligand docking with flexible protein side-chains. Journal of chemical information and modeling, 52(12), pp.3225–3232.

22. Vedamurthy, G.V., Ahmad, H., Onteru, S.K. and Saxena, V.K., 2019. In silico homology modelling and prediction of novel epitopic peptides from P24 protein of Haemonchus contortus. Gene, 703, pp.102–111.

23. Saxena, V.K., Diaz, A. and Scheerlinck, J.P.Y., 2019. Identification and characterization of an M cell marker in nasopharynx-and oropharynx-associated lymphoid tissue of sheep. Veterinary immunology and immunopathology, 208, pp.1–5.

24. Kobylarz, M.J., Grigg, J.C., Liu, Y., Lee, M.S., Heinrichs, D.E. and Murphy, M.E., 2016. Deciphering the substrate specificity of SbnA, the enzyme catalyzing the first step in staphyloferrin B biosynthesis. Biochemistry, 55(6), pp.927–939.

25. Cellini, B., Bertoldi, M., Montioli, R., Laurents, D.V., Paiardini, A. and Voltattorni, C.B., 2006. Dimerization and Folding Processes of Treponema denticola Cystalysin: The Role of Pyridoxal 5 ‘-Phosphate. Biochemistry, 45(47), pp.14140–14154.

26. Livanova, N.B., Chebotareva, N.A., Eronina, T.B. and Kurganov, B.I., 2002. Pyridoxal 5”-Phosphate as a Catalytic and Conformational Cofactor of Muscle Glycogen Phosphorylase b. Biochemistry (Moscow), 67(10), pp.1089–1098.

27. Wardell, S.E., Kwok, S.C., Sherman, L., Hodges, R.S. and Edwards, D.P., 2005. Regulation of the amino-terminal transcription activation domain of progesterone receptor by a cofactor-induced protein folding mechanism. Molecular and Cellular Biology, 25(20), pp.8792–8808..

28. Cai, K., Schirch, D. and Schirch, V., 1995. The affinity of pyridoxal 5′-phosphate for folding intermediates of Escherichia coli serine hydroxymethyltransferase. Journal of Biological Chemistry, 270(33), pp.19294–19299.

29. Deu, E. and Kirsch, J.F., 2007. Cofactor-directed reversible denaturation pathways: the cofactor-stabilized Escherichia coli aspartate aminotransferase homodimer unfolds through a pathway that differs from that of the apoenzyme. Biochemistry, 46(19), pp.5819–5829.

30. Bhatt, A.N. and Bhakuni, V., 2008. Characterization of Pyridoxal 5’-Phosphate-Binding Domain and Folding Intermediate of Bacillus subtilis Serine Hydroxymethyltransferase: an Autonomous Folding Domain. Journal of biochemistry, 144(3), pp.295–303.

31. Akbar, S.M., Sreeramulu, K. and Sharma, H.C., 2016. Tryptophan fluorescence quenching as a binding assay to monitor protein conformation changes in the membrane of intact mitochondria. Journal of bioenergetics and biomembranes, 48(3), pp.241–247.

32. Gaitonde, M.K., 1967. A spectrophotometric method for the direct determination of cysteine in the presence of other naturally occurring amino acids. Biochemical Journal, 104(2), pp.627–633

33. Venkitasubramanian, P., Daniels, L. and Rosazza, J.P., 2007. Reduction of carboxylic acids by Nocardia aldehyde oxidoreductase requires a phosphopantetheinylated enzyme. Journal of Biological Chemistry, 282(1), pp.478–485.

34. Akhtar, M.K. and Jones, P.R., 2014. Cofactor engineering for enhancing the flux of metabolic pathways. Frontiers in bioengineering and biotechnology, 2, p.30.

35. Huang Y, Su L, Wu J (2016) Pyridoxine Supplementation Improves the Activity of Recombinant Glutamate Decarboxylase and the Enzymatic Production of Gama-Aminobutyric Acid. PLoS ONE 11(7): e0157466.

36. Said, Z.M., Subramanian, V.S., Vaziri, N.D. and Said, H.M., 2008. Pyridoxine uptake by colonocytes: a specific and regulated carrier-mediated process. American Journal of Physiology-Cell Physiology, 294(5), pp.C1192–C1197.

37. Schenker, S.T.E.V.E.N., Johnson, R.F., Mahuren, J.D., Henderson, G.I. and Coburn, S.P., 1992. Human placental vitamin B6 (pyridoxal) transport: normal characteristics and effects of ethanol. American Journal of Physiology-Regulatory, Integrative and Comparative Physiology, 262(6), pp.R966–R974.

38. Yamada, R.H., Tsuji, T. and Nose, Y., 1977. Uptake and utilization of vitamin B6 and its phosphate esters by Escherichia coli. Journal of nutritional science and vitaminology, 23(1), pp.7–17.

39. Dempsey, W.B. and Pachler, P.F., 1966. Isolation and characterization of pyridoxine auxotrophs of Escherichia coli. Journal of bacteriology, 91(2), pp.642–645

