## Supplementary figures and images for "Pyridoxal 5’-phosphate supplementation modulates the heterologous expression and activity of a PLP dependent model protein in *E.coli*"

### Fig. A1 Supplementary Figure

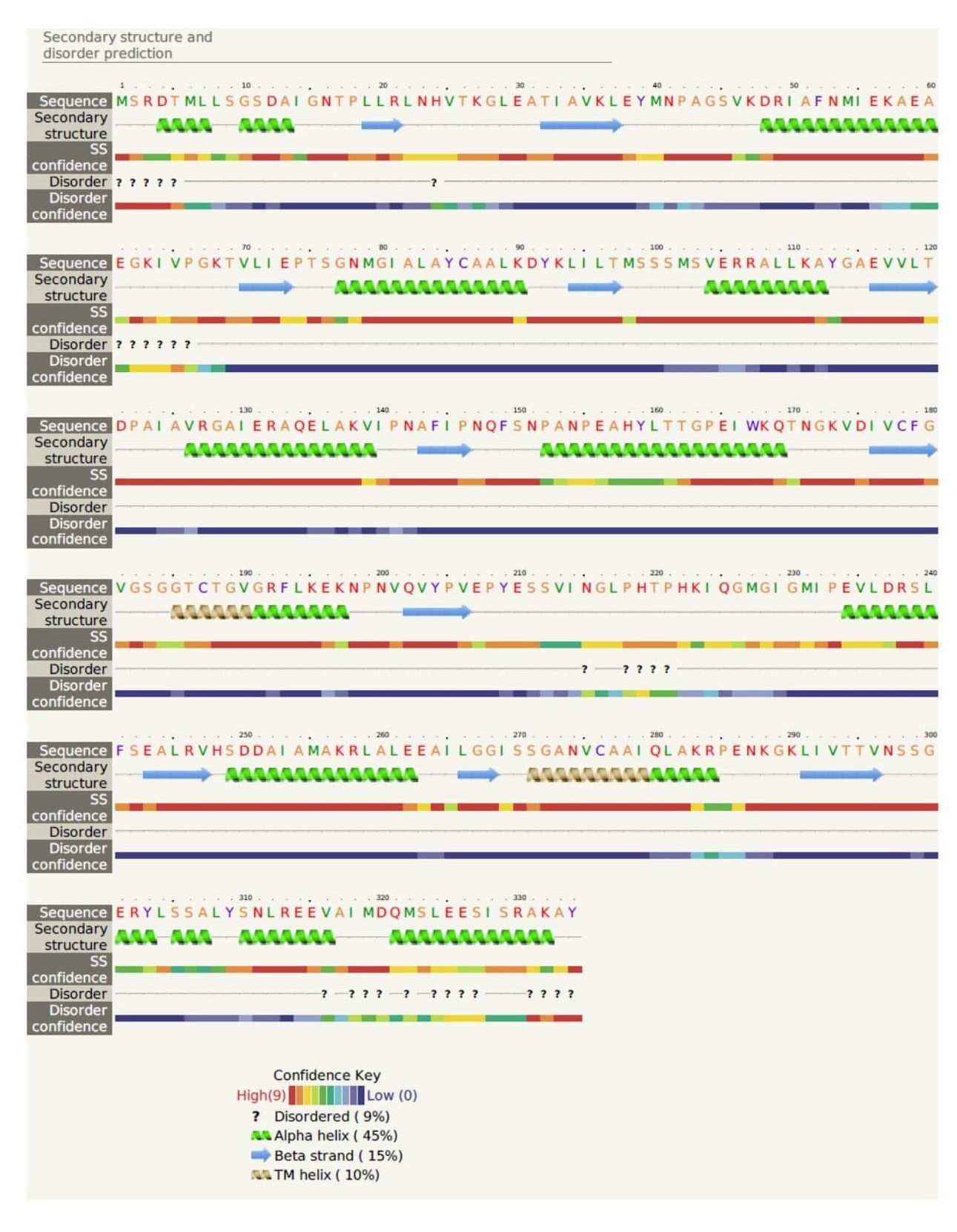

### Fig. A2 Supplementary Figure

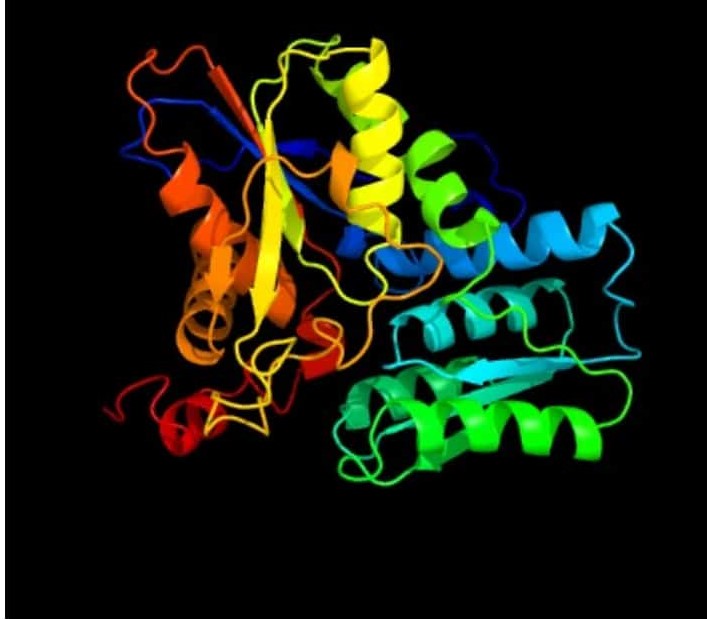

### Fig. A3 Supplementary Figure

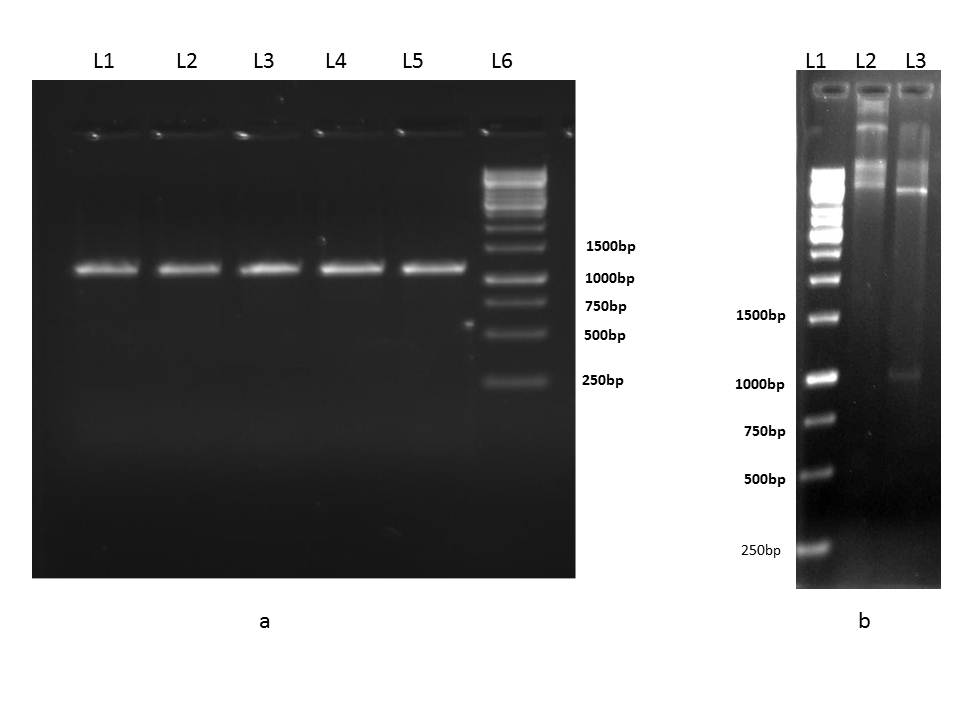
